# A staged evidence chain identifies a recent horizontal transfer of a HET-domain gene into a *Fusarium tricinctum* strain from a lineage related to the *Fusarium oxysporum* species complex

**DOI:** 10.64898/2026.10.12.758604

**Authors:** Sadiya Ahmad, Xiaoqiu Huang

**Affiliations:** Department of Computer Science, Iowa State University, Ames, IA

## Abstract

Genes encoding the heterokaryon incompatibility (HET) domain form fast-evolving, highly paralogous families, which makes horizontal gene transfer (HGT) among them hard to separate from duplication, loss and annotation artefacts. We describe a staged evidence chain for testing a single HGT candidate and apply it to the strongest signal in the HET-domain view of a 109-genome similarity graph of 8,815 HET-domain (PF06985) proteins. Each stage is a separate test with a decision rule fixed before it was run: (i) a gene tree with ultrafast-bootstrap support; (ii) a species-tree check that donor and recipient lie in different clades; (iii) a genome-wide identity background and a contamination screen of the recipient assembly; (iv) a sister-species control that removes genes conserved by vertical descent; and (v) a tblastn search of whole genome assemblies, so that absence is tested in the DNA rather than in the gene annotation. We identify a recent horizontal transfer of a HET-domain gene into one *F. tricinctum* strain (MPI-SDFR-AT-0068) from a lineage related to the *F. oxysporum* species complex (FOSC). The *F. tricinctum* copy is 98.4% identical to its best FOSC hit (local protein identity), against a genome-wide median of 83%, and its synonymous divergence from the FOSC copies (median *d*_*S*_ 0.11) is lower than that of 99.6% of *F. tricinctum* genes from their best *F. oxysporum* hits and of 91% of *F. tricinctum* genes from their best hits in the sister species, which argues strongly against vertical descent, hidden paralogy and shared ancestral alleles. Protein gene trees group it with the FOSC copies (UFBoot 96–98), and a nucleotide tree of the gene region places it as sister to all sampled FOSC copies (UFBoot 98). *F. tricinctum* and the *F. oxysporum* strains belong to different species complexes (species-tree patristic distance 0.075, about 22 times the distance between *F. oxysporum* strains). The gene is absent from the assembly of *F. tricinctum*’s sister species, *F. avenaceum* (best locus 49% identity), and from 14 of 15 further *F. avenaceum* genomes, from 23 of the 26 assemblies of *Fusarium* species outside the FOSC and the *F. tricinctum* complex, and no other sequenced genome of the recipient strain’s own lineage carries it; close copies (≥90%) occur only in 7 of 9 FOSC strains and in one metagenome-assembled *Fusarium* genome from neither complex. In *F. tricinctum* the gene sits in a ∼6-kb strain-specific island on a putative accessory scaffold with *F. oxysporum*-affine gene content, whereas in the FOSC it occupies one conserved, subtelomeric locus of a core chromosome. Assembly contamination was excluded, and single long reads span the gene and its flanks. The donor lineage remains unresolved, and formal reconciliation is still needed. The evidence chain is cheap and uses only public assemblies; each stage targets a specific alternative explanation, and synonymous divergence against the genome background proved the most decisive test.

## 1 Introduction

Heterokaryon incompatibility (HI) is a non-self recognition system of filamentous fungi: when genetically different hyphae of the same species fuse, the resulting heterokaryotic cells are killed or severely inhibited [1, 2, 3]. It is controlled by *het* loci, and many incompatibility systems include a protein with a HET domain (Pfam PF06985), which mediates the cell death [4]. HET-domain genes form large, rapidly diversifying families: in the genomes analysed here a single genome carries up to 499 copies (67–302 per *Fusarium* genome), paralogs within a genome are often very divergent, and the related fungal NOD-like receptors evolve by gene birth and death [5]. Those properties make them interesting candidates for horizontal gene transfer (HGT) and, at the same time, make HGT among them hard to demonstrate. A protein that is unusually similar between two distant taxa can also be explained by differential paralog loss, by a slowly evolving subfamily, by contamination of a draft assembly, or by a missing annotation in the taxa that appear to lack it.

In *Fusarium*, horizontal movement of genetic material is well documented, above all for the accessory chromosomes of the *F. oxysporum* species complex (FOSC; Section 2). Whether HET genes move between *Fusarium* species complexes has not, to our knowledge, been shown. A generic genome-scale HGT screen does not settle the question for a single gene; the candidate has to survive targeted tests that each remove an alternative explanation [7, 8].

Here we set out such a set of tests as an explicit, ordered evidence chain and apply it to the strongest HGT signal among HET-domain genes in a 109-genome panel (in the HET-domain view). The contribution is twofold: (1) a procedure in which every stage has a pre-stated decision rule and a named alternative it is designed to exclude; and (2) a well-supported recent transfer of a HET-domain gene into a *F. tricinctum* strain from a lineage related to the *F. oxysporum* species complex, which we have not found reported before.

## 2 Prior work

### Horizontal transfer from the *F. oxysporum* complex

Ma et al. [6] showed that the lineage-specific chromosomes of *F. oxysporum* f. sp. *lycopersici* are enriched in transposons and effector genes and can be transferred between strains, turning a non-pathogenic strain into a pathogen. Later work confirmed horizontal chromosome transfer within the complex experimentally and in natural populations, including whole pathogenicity chromosomes, in some cases accompanied by whole or partial core chromosomes or rearranged in the recipient [20, 21, 22], and a recent perspective discusses why particular accessory chromosomes are transferred preferentially [23]. Transfers between the *F. oxysporum* complex and other *Fusarium* species are rarer. Van Dam and Rep [24] found mimp transposons and homologs of *SIX* and other effector genes, several of them 97–100% identical in nucleotide sequence to *F. oxysporum* copies, in two *F. hostae* isolates and one *F. proliferatum* isolate, which they considered suggestive of horizontal transfer to or from *F. oxysporum*. Peck et al. [25] identified six putative horizontally transferred regions of 20–200 kb that most likely moved from the *F. oxysporum* complex into populations of *F. xylarioides*, the coffee wilt pathogen; they carry effector genes and transposons, and two of them match parts of the *F. oxysporum* mobile pathogenicity chromosome. In fungi more broadly, giant *Starship* transposons mobilise large cargo regions and, as Gluck-Thaler et al. note, have been implicated in transfers between closely related species [26].

### HET-domain genes and transfer

In *Neurospora crassa*, HET-domain genes are often clustered near telomeres and with lineage-specific genes [27], and in one *F. oxysporum* f. sp. *lycopersici* isolate many lie close to secondary-metabolite and pathogenicity genes, most of them on core chromosomes [28]. *het* loci, including HET-domain genes, are under balancing selection and can show trans-species polymorphism, in which an allele is more similar to one in another species than to alleles of its own species [29, 30, 31]. Huang [28] reported recent horizontal transfer of supernumerary-chromosome regions between distant populations within the *F. oxysporum* complex and proposed that HET-domain genes move together with secondary-metabolite and pathogenicity genes in such transfers; transfer of a HET-domain gene between *Fusarium* species complexes was not examined. Surveys of fungal NOD-like receptors, some of which carry a HET domain, describe their diversity and architecture rather than transfers between species [5].

### The *F. tricinctum* complex

We found no report of horizontal transfer into *F. tricinctum* or *F. avenaceum* from the *F. oxysporum* complex. Published *F. tricinctum* and *F. tricinctum*-complex genomes [33, 34, 35] report assemblies and gene content, not transfers. The two genomes used here come from *Arabidopsis* root isolates sequenced for a study of endophytism [32], which did not examine HGT.

### Search

We searched Europe PMC and the web for combinations of *tricinctum, avenaceum, oxysporum*, “HET domain”, “heterokaryon incompatibility”, “horizontal gene transfer” and “accessory chromosome”. No study reported this gene, this species pair or HGT of a HET-domain gene between *Fusarium* species complexes. Absence from the literature is not proof of priority, and a check by domain experts is still needed.

### Relation to existing methods

The individual tests in our evidence chain are standard. Peck et al. [25] used the same core logic for *F. xylarioides*: identity above the genome-wide divergence background, phylogenetic placement with the donor complex, presence/absence across strains, and genomic location. What we add is (i) decision rules fixed before each test was run, (ii) a sister-species control that removes genes conserved by vertical descent before donor-like genes are counted, and (iii) absence tested in genome assemblies with a per-query self-control rather than in the gene annotation. The chain is a formalisation of current practice for single candidates, not a new kind of inference.

## 3 Results

### 3.1 The evidence chain

Figure 1 and Table 1 summarise the procedure. A candidate enters the chain from a similarity screen: an edge between two proteins from different species whose normalised similarity is unusually high for their taxonomic distance. It then passes through five tests. Each test has a decision rule written down before the test was run, and a failure at any stage stops the chain or forces the conclusion to be reworded. The tests are deliberately ordered from cheap to expensive, and from “is the pattern phylogenetic at all” to “is the absence real”. After the chain had been run, we added analyses that it does not contain: a synonymous-divergence test against the genome background and a nucleotide gene tree (Section 3.8); a search of further genomes of the *F. tricinctum* complex, which also re-checks stage 4 (Sections 3.6 and 3.7); and checks of read support, whole-genome alignments, the FOSC locus and the mobile-element context (Sections 3.5 and 3.7).

**Table 1:**
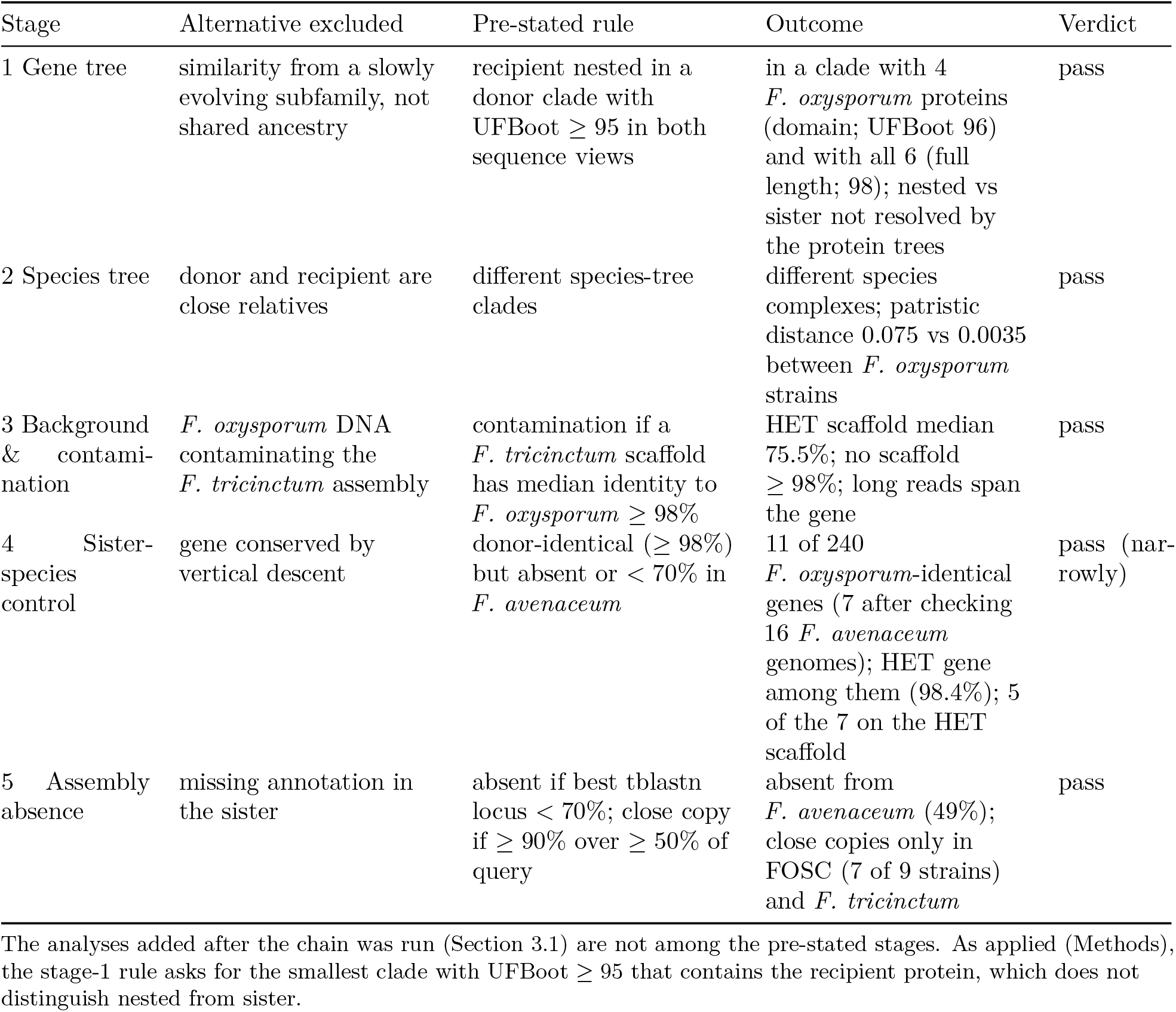
Stages, pre-stated rules and outcomes for the *F. tricinctum* HET candidate.

**Figure 1:**
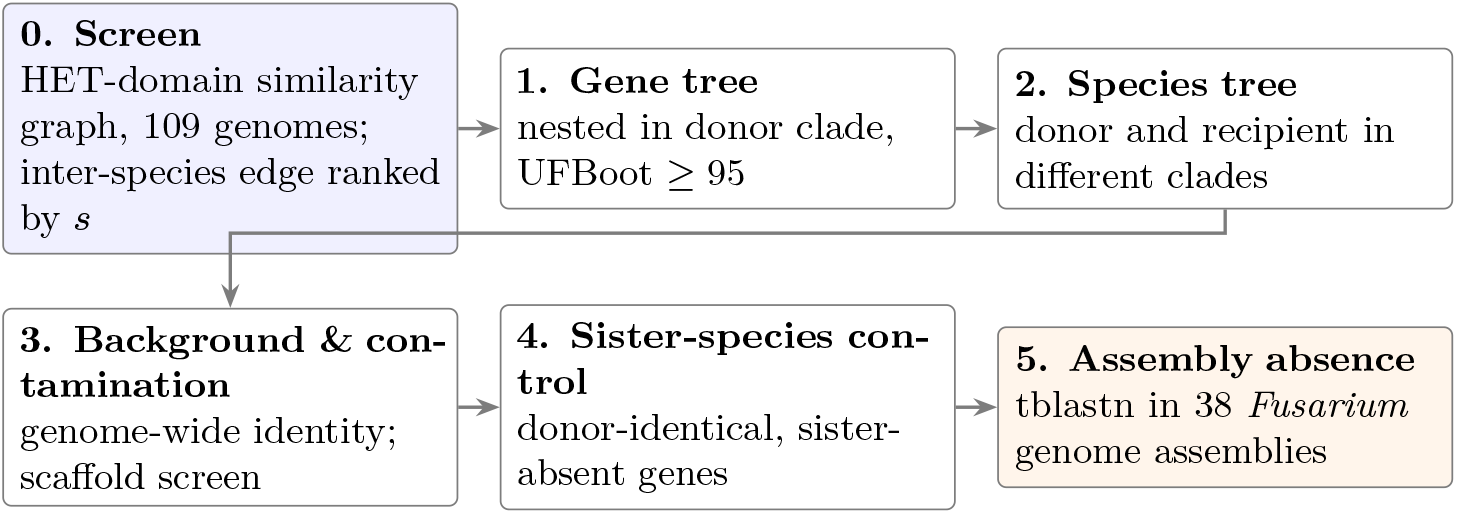
The staged evidence chain. A candidate from the similarity screen (stage 0) is tested in five stages. Every stage has a decision rule fixed before it is run (Table 1) and targets one alternative explanation for the similarity pattern.

### 3.2 Screen: the strongest inter-complex HET signal in *Fusarium*

HET-domain proteins were extracted from 109 fungal proteomes with the PF06985 gathering threshold, giving 8,815 proteins in 89 genomes. All-against-all DIAMOND searches gave a normalised bit-score similarity 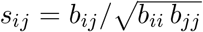 for every pair. Within *Fusarium* (38 genomes, grouped into seven species-tree clusters that largely match the species complexes; the FOSC cluster also contains the *F. fujikuroi* complex and *F. redolens*), the edge with the highest *s* between two different clusters in the HET-domain view joined the *F. tricinctum* protein KAH7230906.1 (gene BKA59DRAFT_488588, assembly GCA_020744515.1) to *F. oxysporum* proteins with identical HET domains (*s* = 1.00, Fo47 QKD61330.2 and f. sp. *lycopersici* 4287 XP_018258429.1): of the 5,470 *Fusarium* HET proteins with at least one inter-cluster edge at *s* ≥ 0.15, it ranked first by raw similarity, tied with these two partners. In the full-length view the signal is weaker; the protein’s best inter-cluster edge (*s* = 0.91) ranks 494th. The *F. tricinctum* protein has no close paralog in its own genome (it forms a singleton within-genome family), so the similarity is not shared with a paralog set. A second *F. tricinctum* HET-domain protein in the screen has a similar profile: KAH7236675.1, on core scaffold 7, is 97.9% identical to an *F. oxysporum* f. sp. *conglutinans* protein, has no *F. avenaceum* hit and forms the highest-scoring *F. tricinctum*–*F. oxysporum* pair in the full-length view (*s* = 0.98). It falls just below the stage-4 cut (98%) and was not taken through the chain here.

### 3.3 Stage 1: the *F. tricinctum* copy groups with the *F. oxysporum* copies in the gene tree

We inferred maximum-likelihood gene trees for the candidate and its 204 closest HET-domain neighbours (up to three per species at *s* ≥ 0.30) in two views: the HET domain aligned to the Pfam HMM, and the full-length protein (Table 2). In both views the *F. tricinctum* protein lies inside a clade of *F. oxysporum* proteins with UFBoot support above the pre-stated threshold of 95: with four *F. oxysporum* proteins in the domain view (UFBoot 96) and with all six annotated *F. oxysporum* copies in the full-length view (UFBoot 98; the seventh, in f. sp. *vasinfectum*, has a truncated gene model that falls below the HET-domain threshold). Its nearest neighbours in the tree are *F. oxysporum* proteins at patristic distances of 0.000–0.032 (domain) and 0.009–0.023 (full length). Whether the *F. tricinctum* copy is nested among the *F. oxysporum* copies or sister to all of them is not resolved by the protein trees. In the domain view its domain is identical to the Fo47 and Fol 4287 domains, so its position among them carries no information. In the full-length tree the split that separates the six *F. oxysporum* copies from it sits on a branch of near-zero length; its UFBoot is 88 in the deposited run and 55–89 in reruns with the same or different settings, and in a re-drawn alignment of the same sequences the *F. tricinctum* copy falls among the *F. oxysporum* copies instead. A nucleotide tree resolves this (Section 3.8). The three *F. tricinctum* paralogs in the tree that are not part of the event fall in unrelated clades: two together with their *F. avenaceum* orthologs (for example KAH7250889.1 with *F. avenaceum* KAH6962732.1, UFBoot 100), and the third (KAH7235560.1) with *F. mangiferae*, because the per-species cap dropped its *F. avenaceum* ortholog. This is as expected under vertical descent.

**Table 2:** Gene-tree placement of *F. tricinctum* KAH7230906.1. IQ-TREE 3.1.4, ModelFinder (BIC), 1,000 UFBoot replicates; supports read per bipartition of the unrooted trees. The “supported clade” is the smallest clade containing the *F. tricinctum* protein with UFBoot ≥ 95.

|  | HET-domain view | Full-length view |
| --- | --- | --- |
| Sequences / alignment columns | 205 / 90 | 205 / 223 |
| Parsimony-informative sites | 78 | 192 |
| Best-fit model | Q.PFAM+R6 | Q.PFAM+R8 |
| Supported clade size | 5 | 7 |
| Clade composition | *F. tricinctum* 1, *F. oxysporum* 4 | *F. tricinctum* 1, *F. oxysporum* 6 |
| Clade UFBoot | 96 | 98 |
| UFBoot for the six *F. oxysporum* copies without *F. tricinctum* | 0 (identical domains) | 88 (near-zero branch) |
| Nearest tips (patristic) | *F. oxysporum* 0.000–0.032 | *F. oxysporum* 0.009–0.023 |

Values are from the deposited run. MAFFT and IQ-TREE run with two threads are not deterministic: in reruns the domain clade has UFBoot 89–96 and the *F. tricinctum* + 6 *F. oxysporum* clade 98–100.

### 3.4 Stage 2: donor and recipient sit in different clades of the species tree

Near-identity matters only if the two species are not close relatives. In the BUSCO-based species tree of the 109 genomes, *F. tricinctum* and *F. avenaceum* are sisters (patristic distance 0.009) inside the *F. tricinctum* species complex, while the nine *F. oxysporum* strains, together with *F. odoratissimum* (also a member of the FOSC), form a tight clade within the cluster that also holds the *F. fujikuroi* complex (Table 3). The two species therefore belong to different species complexes. Both genomes come from the same sequencing project [32]: *F. tricinctum* is strain MPI-SDFR-AT-0068 (GCA_020744515.1), and the *F. avenaceum* comparator is strain MPI-SDFR-AT-0044 (GCA_020744115.1), which was first deposited as *F. tricinctum* and later renamed *F. avenaceum*. The species-tree distance between them (0.0087) is similar to that between *F. graminearum* and *F. culmorum* (0.0092) and larger than several pairs of recognised *F. fujikuroi*-complex species (0.003–0.008). It is consistent with two species, but the naming of both isolates rests on the depositors’ identification. The distance between them is about 22 times the median distance between *F. oxysporum* strains and 8.7 times the distance from *F. tricinctum* to its sister. We note that it lies at only the 30th percentile of the 666 inter-species *Fusarium* distances (median 0.091): the claim is “different species complexes”, not “the most distant parts of the genus”.

**Table 3:** Species-tree patristic distances. (substitutions per site, concatenated BUSCO tree).

| Pair | Distance |
| --- | --- |
| *F. oxysporum* strain vs *F. oxysporum* strain (36 pairs; median, max) | 0.0035, 0.0074 |
| *F. tricinctum* vs *F. avenaceum* (sister species) | 0.0087 |
| *F. tricinctum* vs *F. oxysporum* (9 strains; median, range) | 0.075 (0.075–0.078) |
| All inter-species *Fusarium* pairs (666; median, max) | 0.091, 0.159 |

### 3.5 Stage 3: the signal is not contamination and stands out from the genome background

If the *F. tricinctum* assembly contained *F. oxysporum* DNA, many *F. tricinctum* genes would be near-identical to *F. oxysporum*, and the contaminating scaffolds would be *F. oxysporum*-like as a whole. We searched all 14,106 *F. tricinctum* proteins against nine *F. oxysporum* strains, *F. avenaceum* and *F. graminearum* (best hit with ≥ 50% query coverage). Genome-wide, the median identity of *F. tricinctum* proteins to *F. oxysporum* is 83.0% and to *F. avenaceum* 95.7% (Table 4); the HET gene, at 98.4% to *F. oxysporum* and 49.2% to *F. avenaceum*, is in the bottom 5% of the *F. avenaceum* distribution and in the top 1.3% of the *F. oxysporum* distribution (171 *F. tricinctum* proteins are at least as *F. oxysporum*-identical, most of them genes conserved across *Fusarium*; Section 3.6). The scaffold that carries it (JAGPXF010000009.1, 1.29 Mb, 339 genes) has a median identity to *F. oxysporum* of 75.5%, below the genome-wide value, and no *F. tricinctum* scaffold reaches the pre-stated contamination threshold of a median ≥ 98% to *F. oxysporum*. Long reads confirm this. PacBio reads of the sequenced strain cover scaffold 9 at the same median depth as the core scaffolds (25*×* vs 25–26*×*), six single reads span the whole HET gene with at least 0.5 kb of scaffold-9 sequence on both sides, and BUSCO duplication is 0.7%, so the assembly does not contain a second genome. Contamination is excluded.

**Table 4:** Genome-wide background, the HET scaffold, and the eight core scaffolds of *F. tricinctum*. Best-hit identity, DIAMOND `–sensitive`, ≥ 50% query coverage. “*F. oxysporum*-affine”: ≥ 90% to *F. oxysporum* and ≥ 10 points closer to *F. oxysporum* than to *F. avenaceum* (genes without an *F. avenaceum* hit count as closer).

|  | HET gene | Scaffold 9 (339 genes) | Core scaffolds 1–8 (13,747 genes) |
| --- | --- | --- | --- |
| Identity to *F. oxysporum* (median) | 98.4 | 75.5 | 83.2 |
| Identity to *F. avenaceum* (median) | 49.2 | 50.9 | 95.8 |
| No *F. avenaceum* hit<sup>a</sup> | – | 40.7% | 6.3% |
| *F. oxysporum*-affine genes | yes | 15.9% | 0.6% |
| Best Fol4287 hit on lineage-specific chr. 3/6/14/15<sup>b</sup> | – | 39% | 1.8% |
| Genes $\geq 98\%$ to *F. oxysporum* | yes | 1.5% | 1.8% (genome-wide) |

<sup>a</sup>One search against the combined proteomes with up to 50 targets per query; a separate search against *F. avenaceum* alone gives 35.7% and 6.1%. <sup>b</sup>Share of the 270 (scaffold 9) and 12,295 (core) genes with a Fol 4287 hit. Several genes have equally good hits on different chromosomes, so the value depends on tie-breaking: the pipeline’s sorting rule gives 41% and 2.0%, and other tie-breaks give 34–43% and 1.6–3.8%.

The same comparison shows that scaffold 9 is unusual. The *F. tricinctum* assembly is close to chromosome level, with eight large core scaffolds; scaffold 9 is smaller and its gene content is poorly shared with the sister species (41% of its genes have no *F. avenaceum* hit, against 6% on the core).

Sixteen percent of its genes are *F. oxysporum*-affine, spread along the whole scaffold (Fig. 2), and 39% of those with an *F. oxysporum* f. sp. *lycopersici* 4287 hit have their best hit on that strain’s lineage-specific chromosomes [6] (Table 4, note b). Whole-genome alignment agrees: scaffold 9 aligns to the *F. avenaceum* assembly over only 11% of its length (core scaffolds 82%), about half of it is present in some other genomes of the *F. tricinctum* complex and absent from others, and blocks of it are ≥ 95% identical to *F. oxysporum* accessory chromosomes (Fo47 chromosome VII; Fol 4287 lineage-specific chromosomes 3 and 14). These are the signatures of an accessory chromosome that shares content with *F. oxysporum* accessory chromosomes.

**Figure 2:**
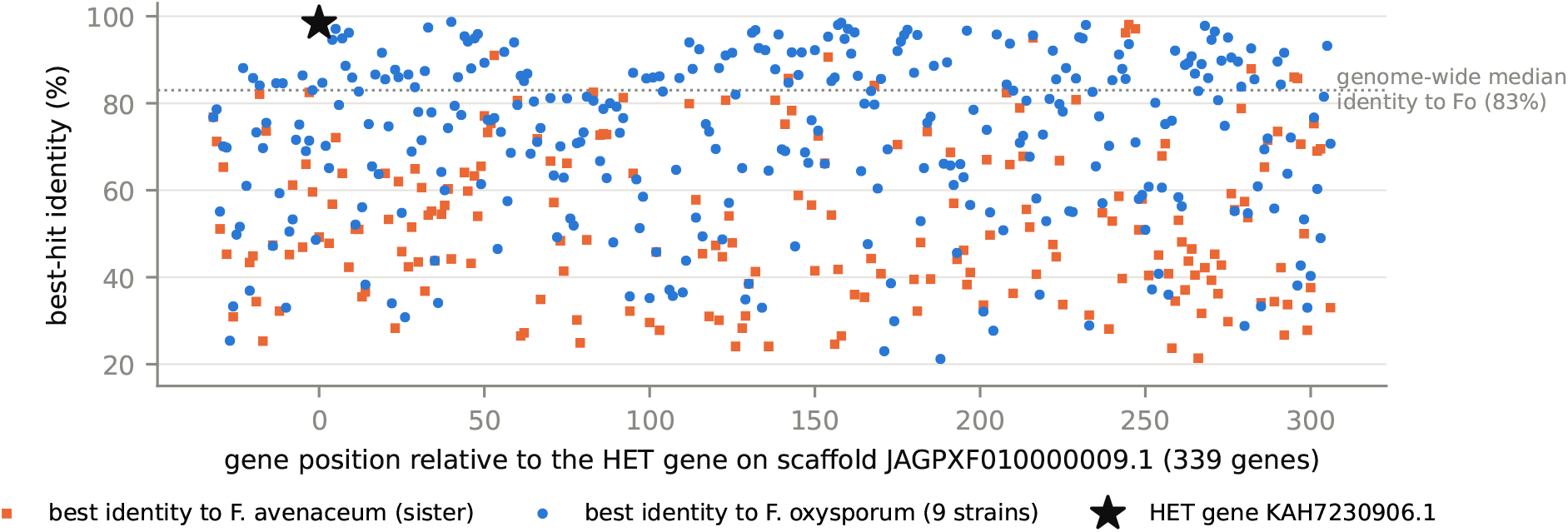
Identity profile of the *F. tricinctum* HET scaffold. For each gene on scaffold JAG-PXF010000009.1, best-hit identity to *F. oxysporum* (blue circles, nine strains) and to the sister species *F. avenaceum* (orange squares). The HET gene (star) is among the most *F. oxysporum*-like genes on the scaffold (two other genes reach 98.7 and 98.5%) and has no close *F. avenaceum* match. The scaffold as a whole is not *F. oxysporum*-like (median 75.5%, dotted line shows the genome-wide median of 83%), which argues against contamination, but *F. oxysporum*-affine genes occur along its whole length.

Only a gene-sized segment is shared with the *F. oxysporum* copies at ≥ 95% nucleotide identity: 2.1 kb (the coding sequence with about 0.2 kb on each side; 96–97% identity) with six of them and 2.8 kb with the f. sp. *cubense* copy. Beyond it, the flanking sequences of the *F. tricinctum* and *F. oxysporum* copies are unrelated. In MPI-SDFR-AT-0068 the gene lies in a ∼6-kb island that the other genomes of its lineage lack (Section 3.7). The island is bounded by two inverted, 81%-identical copies (1.7 and 2.2 kb) of an unclassified repeat and contains no transposon-related open reading frame (a Tc5-type transposase-domain ORF lies 0.5 kb outside it); the nearest *Starship* captain gene lies 34.6 kb away and has no close homolog in *F. oxysporum* (≤ 40% identity).

### 3.6 Stage 4: a sister-species control isolates donor-like genes

High identity to *F. oxysporum* alone is not informative, because genes conserved across *Fusarium* can be near-identical by vertical descent. Of the 240 *F. tricinctum* proteins whose best *F. oxysporum* hit is ≥ 98% identical, 228 are also ≥ 95% identical to *F. avenaceum*, i.e. conserved genes. Only 11 are *F. oxysporum*-identical while absent from, or < 70% identical to, *F. avenaceum* (Table 5). With a single *F. avenaceum* genome, however, “absent from *F. avenaceum*” can be a strain-level or annotation-level absence. We therefore checked the 11 genes in 15 further *F. avenaceum* genomes, and with spliced alignment in the panel’s own *F. avenaceum* assembly. Four have close copies (≥ 90%) in *F. avenaceum*: KAH7241223.1 and KAH7241224.1 in *F. avenaceum* NC74 (96.9% and 97.6% by spliced alignment), KAH7235317.1 in *F. avenaceum* F156N33 (91.8% by tblastn), and KAH7257287.1, which is present, unannotated, in the panel *F. avenaceum* assembly. Seven genes remain donor-like. The HET gene is one of them, and 5 of the 7 lie on scaffold 9, which carries only 2.4% of the genome’s genes. Applied literally to all 16 *F. avenaceum* genomes, the stage-4 rule (< 70%) would remove three more, because some *F. avenaceum* genomes carry related copies of KAH7231063.1 and KAH7231064.1 (80–87%) and one (*F. avenaceum* XZ-22-127) carries an 82.7% copy related to the HET gene. For the HET gene that copy belongs to a separate subfamily (Section 3.8); for the other two this has not been tested. The donor-like signal is therefore concentrated on the HET scaffold but dispersed along it: several *F. oxysporum*-like pieces on an accessory-like scaffold, each of which may have its own history. The HET gene is not the most recent of them: two adjacent scaffold-9 genes, KAH7231063.1 and KAH7231064.1, are 4.6 and 6.5 times less diverged from their best *F. oxysporum* hits at synonymous sites (*d*_*S*_ 0.020 and 0.014, against 0.089 for the HET gene).

**Table 5:**
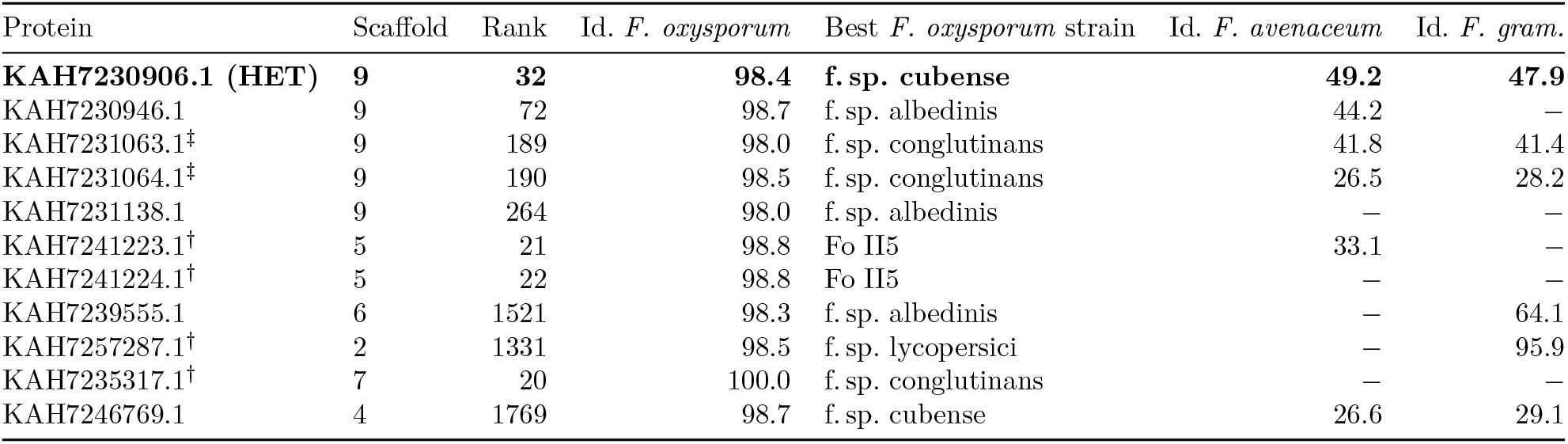
The 11 *F. tricinctum* proteins whose best *F. oxysporum* hit is ≥ 98% identical but that are absent (*−*) or < 70% identical in the panel *F. avenaceum* genome. Rank = gene order on the scaffold, counted from 0. Scaffold 9 is the HET scaffold. *†*Close copy (≥90%) in another *F. avenaceum* genome or, unannotated, in the panel *F. avenaceum* assembly; not donor-like. ‡Related copies (80–87%) in some other *F. avenaceum* genomes, whose relationship to these genes is untested (see text).

The HET gene passes this stage narrowly. Its 98.4% is the local identity over residues 1–506 to its best-matching strain (*F. oxysporum* f. sp. *cubense*); against Fo47, Fol 4287 or the *radicis-cucumerinum* strain alone it is 97.4–97.8%, and at a 99% cut it would not pass. Its absence from the sister species, by contrast, is robust (Section 3.7).

### 3.7 Stage 5: the gene is absent from the sister species’ genome assembly

Absence from a proteome can mean the gene was not annotated. We therefore searched the genome assemblies of all 38 *Fusarium* genomes in the panel with tblastn, chaining HSPs on the same contig and strand into loci so that introns do not split a gene. The rules were fixed before the search: a *close copy* is a locus with ≥ 90% identity over ≥ 50% of the query; *absent* means the best locus is < 70% identical (HET paralogs at 40–60% exist in every *Fusarium* genome); anything in between is a *related copy*. A self-search of each query against its own *F. tricinctum* assembly served as a positive control.

For the HET gene (Fig. 3, Table 6), the self-control recovered the gene at 99.6% over the full query. Close copies (96.7–97.7%) are present in 7 of the 9 *F. oxysporum* strains and nowhere else. The best locus in the *F. avenaceum* assembly is 49% identical, a HET paralog, so the gene is missing from the sister species’ DNA and not just from its annotation. The 23 other assemblies outside the FOSC in which it is absent, including all *F. solani*-complex genomes and most of the *F. fujikuroi* and *F. sambucinum* complexes, have best loci of 41–50%; it is also absent (42%) from the *F. odoratissimum* assembly, a FOSC member that is the same strain as *F. oxysporum* II5. Related copies at 80–82% occur in *F. oxysporum* f. sp. *conglutinans, F. proliferatum, F. culmorum* and *F. flagelliforme*; two of them (*F. proliferatum, F. flagelliforme*) belong to the related subfamily of Fig. 6. Within the FOSC the gene sits at one orthologous, subtelomeric locus. Five of the seven chromosomes or scaffolds that carry the close copies align mainly (73–82%) to Fol 4287 chromosome 12, a core chromosome; the f. sp. *cubense* copy lies on a mosaic scaffold, and Fol 4287’s own copy on an unplaced scaffold. At least one 30-kb flank of every copy maps to that unplaced Fol 4287 scaffold, and in five of the seven assemblies the copy lies 68–363 kb from a contig end. The two strains without a close copy (Fo II5, which has none, and f. sp. *conglutinans*, which has only a related copy) therefore reflect variation at this subtelomeric locus, not presence or absence of an accessory chromosome.

**Table 6:** tblastn absence check of the HET gene and the four other scaffold-9 genes from Table 5 across 38 *Fusarium* assemblies. Identities are of the best chained locus.

| Query | Self | *F. avenaceum* | Close / related / absent | Close copies in |
| --- | --- | --- | --- | --- |
| **KAH7230906.1 (HET)** | **99.6** | **49.2** | **8 / 4 / 26** | ***F. oxysporum* (7 of 9 strains), *F. tricinctum*** |
| KAH7231063.1 | 100.0 | 38.7 | 2 / 2 / 34 | *F. oxysporum* f.sp. *conglutinans*, *F. tricinctum* |
| KAH7231064.1 | 98.4 | 28.5 | 2 / 2 / 34 | *F. oxysporum* f.sp. *conglutinans*, *F. tricinctum* |
| KAH7231138.1 | 100.0 | no hit | 14 / 1 / 23 | 10 FOSC assemblies (9 strains; incl. *F. odoratissimum*), 2 *F. solani*-complex, *F. redolens*, *F. tricinctum* |
| KAH7230946.1* | 82.4 | 44.2 | 1 / 4 / 33 | *F. oxysporum* f.sp. *albedinis* |

\*Self-control below the close-copy threshold (82% to its own assembly) because tblastn HSPs run into short introns; the row is shown for completeness.

**Figure 3:**
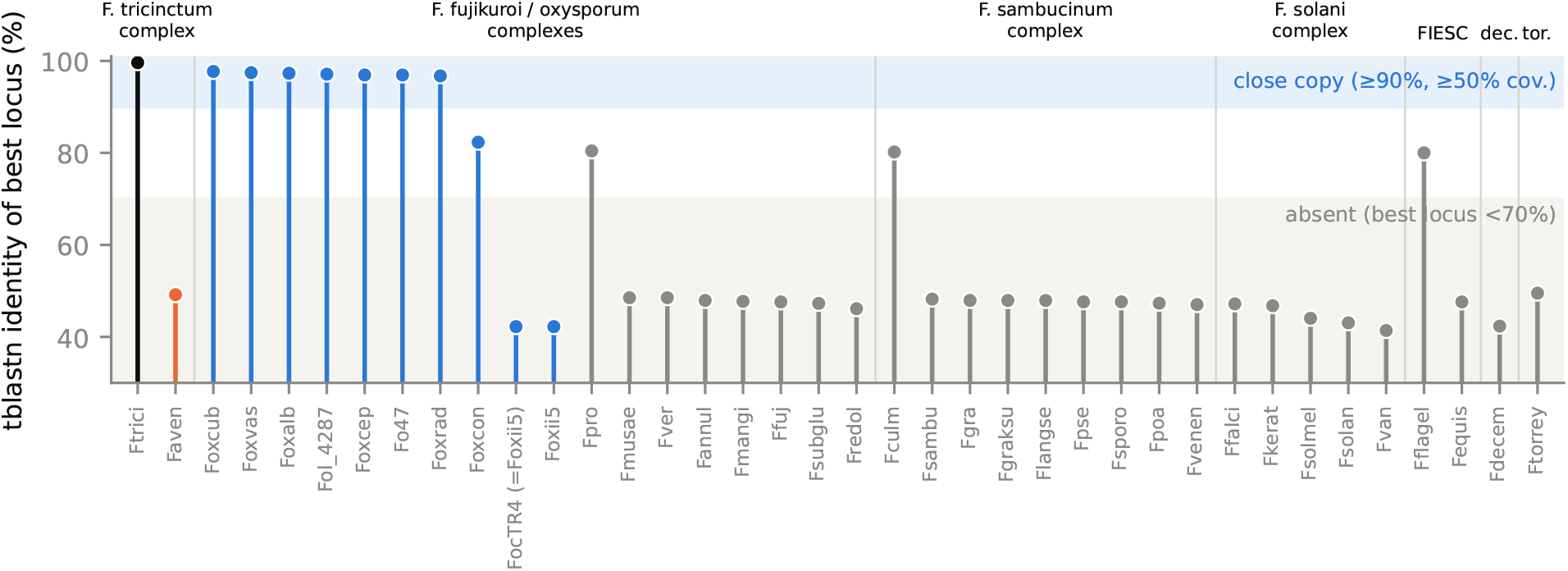
tblastn search for the HET gene in 38 *Fusarium* genome assemblies. Each stem is the identity of the best chained locus in one assembly, grouped by species-tree cluster (the *F. fujikuroi*/*oxysporum* cluster also contains *F. redolens*). Black: *F. tricinctum* self-control; orange: the sister species *F. avenaceum*; blue: FOSC strains, including *F. odoratissimum* (FocTR4), which is the same strain as *F. oxysporum* II5 (Foxii5); grey: other species. Shaded bands mark the pre-stated calls: close copy (≥ 90% identity over ≥ 50% of the query) and absent (< 70%). FIESC, *F. incarnatum-equiseti* complex; dec., *F. decemcellulare*; tor., *F. torreyae*.

All four other scaffold-9 genes are also absent from the *F. avenaceum* assembly (best loci 28–44%, or no hit). Two adjacent genes, KAH7231063.1 and KAH7231064.1, have close copies only in *F. oxysporum* f. sp. *conglutinans* among the panel genomes (the same adjacent pair is also present, at 87–90%, in *F. subglutinans*), which suggests that the *F. oxysporum*-like material on scaffold 9 did not all come from the same *F. oxysporum* lineage, or from one lineage that was later reshuffled among *F. oxysporum* strains. KAH7231138.1 has a wider, patchy distribution across three species complexes, so it is not evidence for an *F. oxysporum*-specific source.

#### Further genomes of the *F. tricinctum* complex

To test whether the gene is a property of the species or of one strain, we searched 25 further public assemblies labelled as species of the *F. tricinctum* complex with the same procedure: six *F. tricinctum* (five strains), 15 *F. avenaceum*, three “*F. acuminatum*” and one *F. flocciferum* (Fig. 4). One of the “*F. acuminatum*” assemblies is a metagenome-assembled genome from another species complex (see below). The gene is absent from all six *F. tricinctum*-labelled genomes and from 14 of the 15 *F. avenaceum* genomes (best loci 46–49%, HET paralogs). Four genomes carry related copies of 76–83% (*F. avenaceum* XZ-22-127, *F. flocciferum* and two “*F. acuminatum*” assemblies); the three in the gene trees (all but the 76% fragment in F829) belong to a separate subfamily (Section 3.8). None of the four other sequenced genomes of the recipient’s own lineage, which are ∼99% identical to it genome-wide, carries the gene: *F. tricinctum* MES5 and the gapless *F. tricinctum* MsR-QD66 lack it, and two assemblies labelled *F. acuminatum* (F829 and 1A) carry only a related copy. In MsR-QD66 the flanking genes are present on chromosome 10, but the ∼6-kb window that holds the gene in MPI-SDFR-AT-0068 is not. The transfer is therefore specific to strain MPI-SDFR-AT-0068: the gene was acquired after this lineage diversified, or lost from its relatives. Outside the panel, one close copy (95.6% over the full length) occurs in a public metagenome-assembled genome (MAG; GCA_027247665.1, labelled “*F. acuminatum*” isolate CHS3, from a root metagenome), on a 5-Mb contig. That genome is not a member of the *F. tricinctum* complex: genome-wide it is closest to the *F. sambucinum* complex (∼88% nucleotide identity).

**Figure 4:**
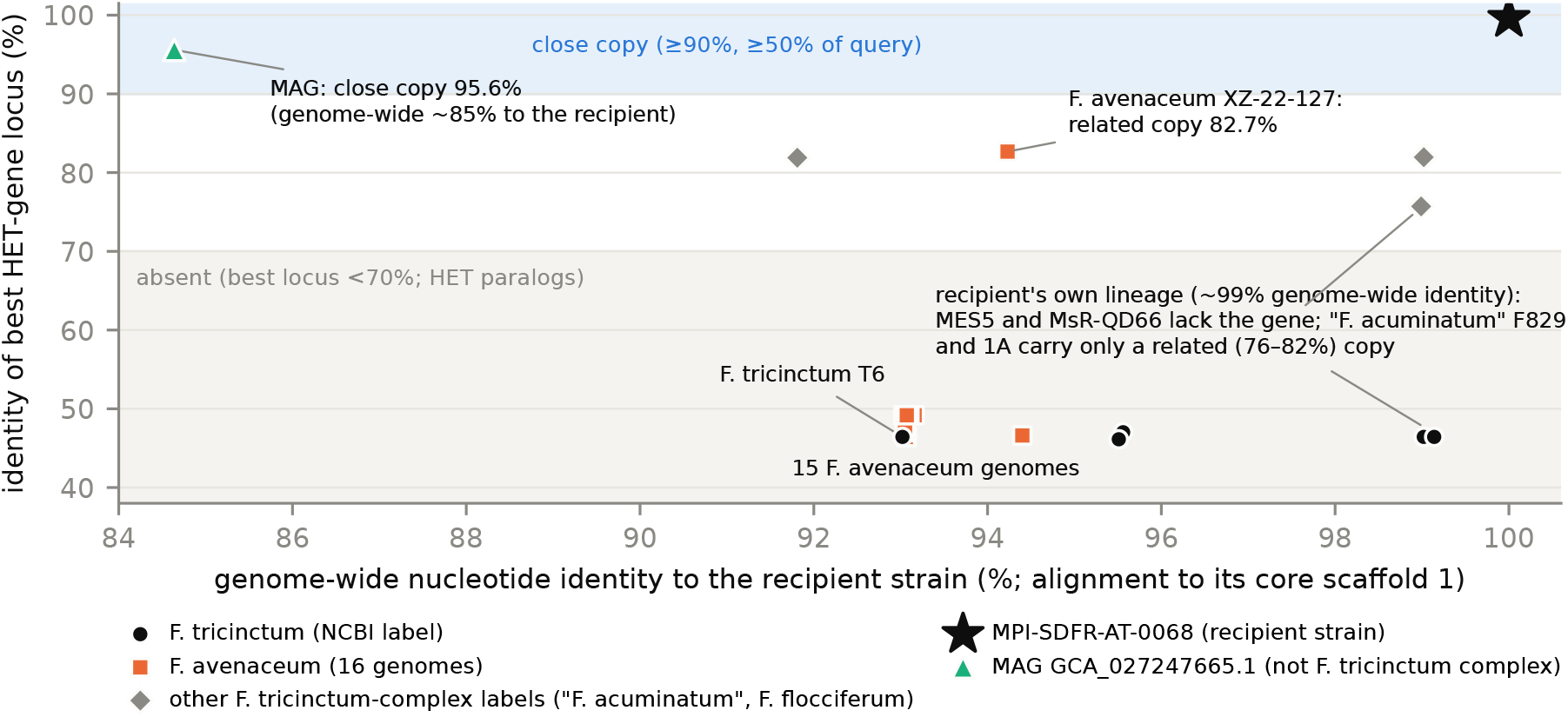
The HET gene in 26 assemblies (25 strains) of the *F. tricinctum* species complex and one metagenome-assembled genome. For each assembly, the identity of the best tblastn locus for the HET gene (same chaining and calls as Fig. 3) is plotted against its genome-wide nucleotide identity to the recipient strain (minimap2 alignment to its 7.2-Mb core scaffold 1). Genomes ∼99% identical to the recipient (its own lineage) lack a close copy, so the gene is strain-specific. The only close copy outside the recipient and the FOSC is in the MAG GCA_027247665.1, which is not a member of the *F. tricinctum* complex. Overlapping points: 11 *F. avenaceum* genomes coincide near 93%/49%, and three more lie under the T6 point. The MAG’s genome-wide identity rests on the 16% of contig 1 that aligns to it.

### 3.8 Direction of transfer and the donor

Two observations point to transfer into this *F. tricinctum* strain rather than out of it. First, the gene is absent from *F. tricinctum*’s sister species (in 15 of 16 genomes), and no other sequenced genome of the strain’s own lineage carries it, so the *F. tricinctum* copy was not inherited from the *F. tricinctum*–*F. avenaceum* ancestor; within the strain’s lineage it was either acquired after the lineage diversified or lost from the strain’s relatives. Second, the genomic context differs between the two sides. In the FOSC the gene occupies one conserved, subtelomeric locus, shared by seven strains, which indicates long-term residence. In *F. tricinctum* it sits in a ∼6-kb strain-specific island on an accessory-like scaffold, without *F. oxysporum* flanking DNA (Sections 3.5 and 3.7).

Synonymous divergence dates the event against the genome background (Fig. 5). Between the *F. tricinctum* copy and the six annotated *F. oxysporum* copies, *d*_*S*_ is 0.09–0.12 (Nei–Gojobori; median 0.106). The median is lower than the *d*_*S*_ of 99.6% of 13,152 *F. tricinctum* genes to their best *F. oxysporum* hits (median 1.17 among the 85% of pairs that are not saturated), lower than that of 97% of the 228 conserved genes that are ≥ 98% identical to *F. oxysporum* in protein sequence (median 0.52), and lower than that of 91% of *F. tricinctum* genes to their best *F. avenaceum* hits (median 0.19). Other estimators, reciprocal-best-hit backgrounds and the in-frame part of the gene give the same picture (Methods). Under vertical descent from the ancestor of the two species complexes, the copies would have diverged like the rest of the genome. Instead, the *F. tricinctum* and *F. oxysporum* copies share a common ancestor far more recently than the two complexes split and, if synonymous rates are similar across genes, more recently than *F. tricinctum* and *F. avenaceum* split.

**Figure 5:**
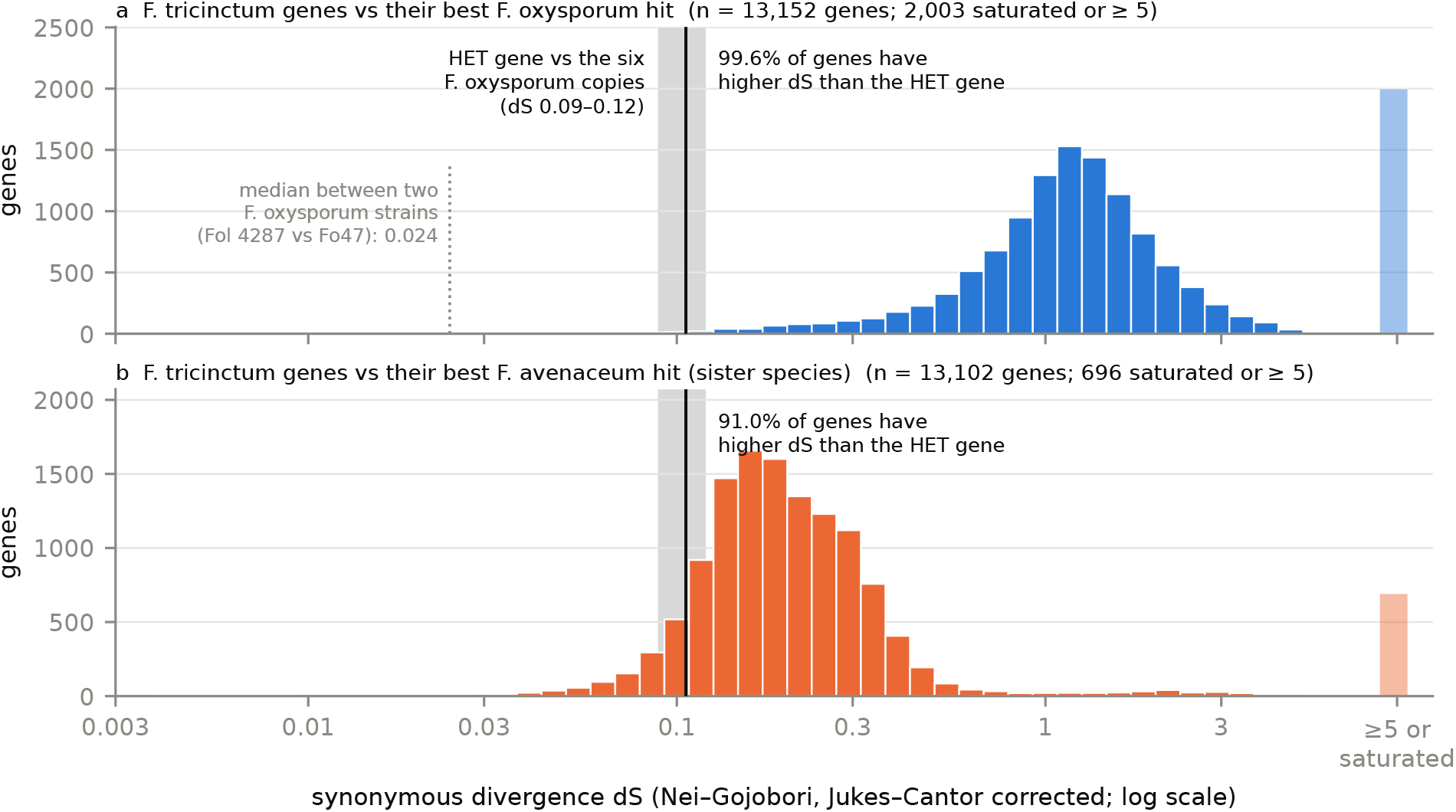
Synonymous divergence of the HET gene against the genome background. *d*_*S*_ (Nei– Gojobori, Jukes–Cantor corrected) of the 13,152 (a) and 13,102 (b) *F. tricinctum* genes that have a best hit in nine *F. oxysporum* strains (a) or in the sister species *F. avenaceum* (b) with more than 50 aligned codons; pairs with saturated synonymous sites (*p*_*S*_ ≥ 0.75) or *d*_*S*_ ≥ 5 are pooled at the right, and values below 0.003 (including 0) at the left. The grey band and black line show the HET gene against the six annotated *F. oxysporum* copies (*d*_*S*_ 0.089–0.120; median 0.106). The dotted line in (a) is the median *d*_*S*_ between two *F. oxysporum* strains (Fol 4287 vs Fo47; best hits of a random sample of 1,500 Fol 4287 proteins; unsaturated pairs).

The donor is less clear. A nucleotide tree of the gene region places the *F. tricinctum* copy as sister to all sampled FOSC copies (Fig. 6): the seven FOSC copies form a clade that excludes it (SH-aLRT 96–97, UFBoot 98, depending on the copies included), and at least six sites carry a base shared by all FOSC copies while the *F. tricinctum* copy keeps the outgroup state. In the 11-sequence analysis, AU tests reject 12 of 14 constrained trees that place the *F. tricinctum* copy inside the FOSC clade, including its grouping with any single FOSC copy (p ≤ 0.045; p = 0.03 for the f. sp. *cubense* copy). The two constrained trees that are not rejected (p ≈ 0.07) force it into a clade with six of the seven copies, and their best trees place it next to the *cubense* copy or next to five copies. With the related copies added, 10 of the 14 are rejected, and a sister relation to the *cubense* copy is not (p = 0.06). Synonymous divergence does not settle this: on the in-frame part of the gene, *d*_*S*_ between the *F. tricinctum* copy and the FOSC copies (0.067–0.098, median 0.086) overlaps that among the FOSC copies themselves (0.005–0.086, median 0.049). The best-supported reading is a donor lineage that diverged before the sampled FOSC copies did, either an unsampled or early-diverging FOSC lineage or a third lineage; a source close to the least-derived sampled copy (f. sp. *cubense*) is not excluded. A third lineage is a real possibility: the MAG copy (Section 3.7) is the closest relative of the *F. tricinctum* + FOSC clade, and grouping it with the *F. tricinctum* copy is not rejected in the protein tree (p = 0.06–0.12) or in the 11-sequence nucleotide tree (p = 0.07), although it is in the 16-sequence tree (p = 0.04). The direction of transfer, into *F. tricinctum*, is well supported; the donor lineage is not resolved.

**Figure 6:**
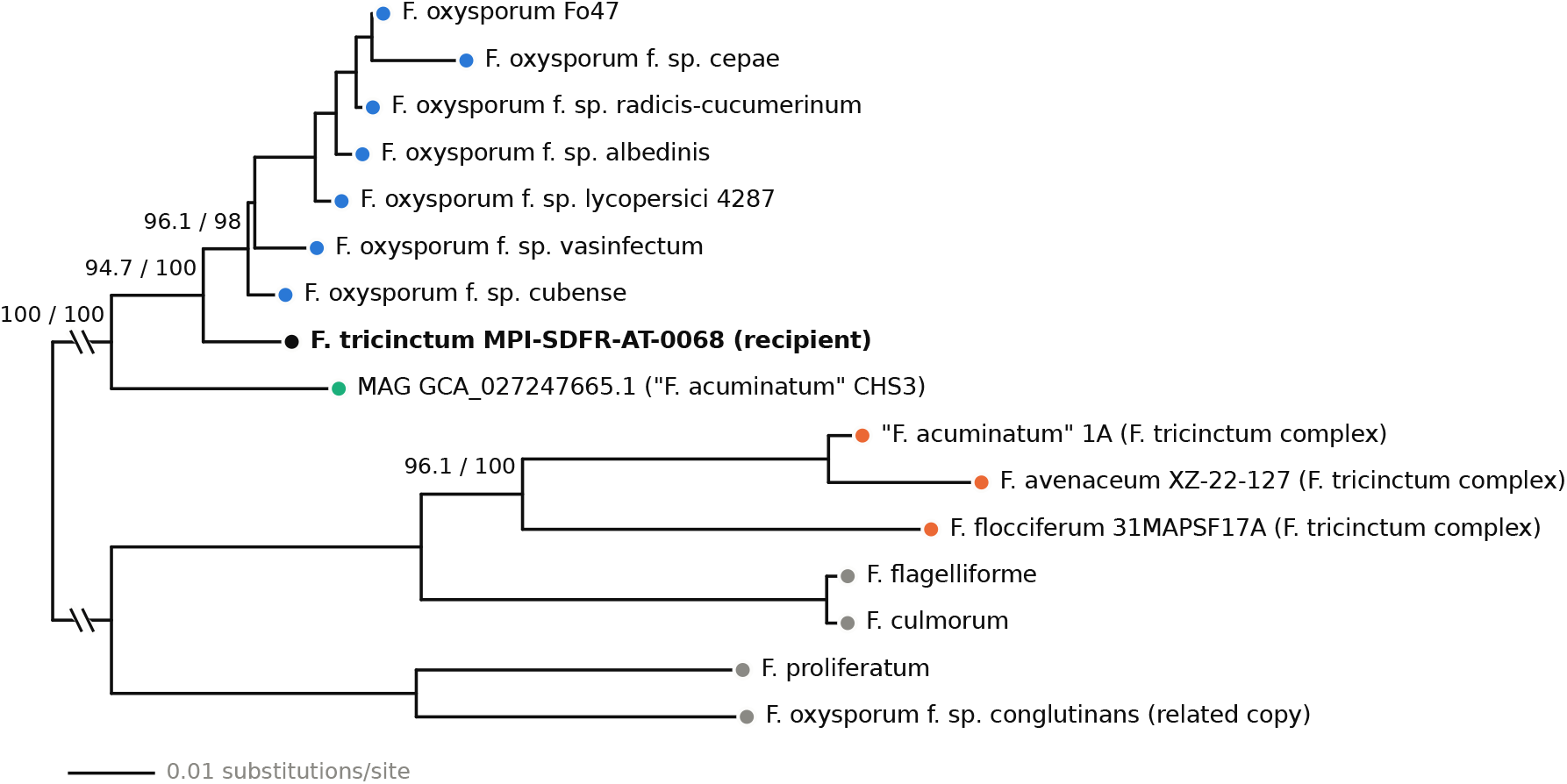
Nucleotide gene-region tree of the HET gene and its related copies. Maximum-likelihood tree (IQ-TREE 3.1.4, K2P+I; 16 sequences, 1,629 alignment columns covering the region homologous to the *F. tricinctum* gene, intron included) of the *F. tricinctum* copy (black), the seven FOSC copies (blue; all copies were located by spliced alignment of the *F. tricinctum* protein, so f. sp. *vasinfectum*, whose annotated gene model is truncated, is included), the MAG copy (green), the related (∼80–82%) copies from three *F. tricinctum*-complex genomes (orange; the partial copy in F829 is not included) and from other *Fusarium* species, including *F. oxysporum* f. sp. *conglutinans* (grey). Numbers are SH-aLRT / UFBoot for the splits discussed in the text, read per bipartition. The tree is rooted between the close copies and the related subfamily; that stem (0.13 substitutions/site) is drawn shortened. With only *F. proliferatum* and *F. flagelliforme* as outgroups (11 sequences, HKY+G4) the topology of the close copies is the same (FOSC clade 97 / 98).

## 4 Discussion

The procedure turns a similarity outlier into a defensible HGT candidate by testing alternatives one at a time. The gene tree addresses “a slowly evolving subfamily”; the species tree, “close relatives”; the genome background and scaffold screen, contamination; the sister-species control, vertical conservation; and the assembly search, annotation gaps. Applying it showed where the chain needs strengthening. Synonymous divergence against the genome background separates transfer from vertical descent and from old shared alleles more directly than protein identity or tree nesting can. A nucleotide tree was needed because the protein trees could not resolve whether the recipient copy is nested among or sister to the donor-complex copies. And the sister-species control needs more than one sister genome: with one, 4 of its 11 calls were strain-level or annotation-level absences. Every stage of the chain used a decision rule written down before it was run (the stage-1 protocol, i.e. neighbour threshold, alignment and trimming, was revised after a first run, without changing this candidate’s placement in the domain view; Methods); the analyses added afterwards had no pre-stated rules.

The candidate sits on a scaffold with the hallmarks of an accessory chromosome and with dispersed *F. oxysporum*-affine content, which suggests that this *F. tricinctum* lineage has exchanged accessory material with relatives of the FOSC. The HET gene itself, however, did not arrive as part of such a block: only a gene-sized segment (2–3 kb) is shared with the *F. oxysporum* copies, it sits in a ∼6-kb island that only this strain carries, and in the FOSC it resides at a conserved, subtelomeric locus of a core chromosome. Its transfer looks like a separate, gene-sized event into an accessory-like region. Because the HET-domain family includes non-self recognition genes, the acquisition of a near-identical HET-domain gene from a relative of the FOSC raises the question of whether such transfers change compatibility interactions between species that share a host; that question needs experimental work.

### Trans-species polymorphism as an alternative

*het* loci are under balancing selection, and old allele classes can be shared across species [29, 30, 31]. Under trans-species polymorphism, however, the shared alleles would have diverged since before the species split, so their divergence would be at or above the genome background (at the HET-domain locus *het-e* of *N. crassa*, three times the genome average [30]). The *F. tricinctum* and *F. oxysporum* copies show the opposite: their median *d*_*S*_ is lower than that of 99.6% of genes between the two genomes and of 91% of genes between *F. tricinctum* and its sister species (Section 3.8). Related (∼82%) copies do occur in other genomes of the *F. tricinctum* complex (Figs 4 and 6), but they belong to a separate subfamily, and the *F. tricinctum* copy does not group with them (AU p < 0.001 in both the protein and the nucleotide tree). We therefore consider trans-species polymorphism very unlikely; a population sample of *F. avenaceum* and *F. tricinctum* would test it further.

### Comparison with the *F. xylarioides* case

The scaffold-level pattern resembles the *F. oxysporum → F. xylarioides* transfers [25]: *F. oxysporum*-like material scattered on a recipient region, and patchy presence among *F. oxysporum* strains that differ in which strain is the closest match (here, f. sp. *conglutinans* for two neighbouring scaffold-9 genes). The HET gene differs in mechanism: a gene-sized unit moved, not a 20–200-kb region, and no *Starship* or recognisable transposon lies within the island that carries it. Unlike that case, the transferred gene is a HET-domain gene rather than an effector.

### Limitations

(1) The event is strain-specific: no other sequenced genome of the recipient’s lineage carries the gene (Section 3.7; the *F. tricinctum*-complex hazelnut isolate of Turco et al. [34] was not searched). Whether this reflects a very recent acquisition or loss in the other strains, and where the donor lineage sits, needs more genomes from the FOSC, the *F. tricinctum* complex and the *F. sambucinum* complex. (2) Absence from *F. avenaceum* rests on assemblies. The panel *F. avenaceum* genome is a 13-contig PacBio assembly, and the gene is also absent from 14 of 15 further *F. avenaceum* genomes, so an assembly gap is unlikely. For the other sister-absent genes one sister genome was not enough (Section 3.6). (3) No formal gene-tree/species-tree reconciliation (e.g. AleRax) has been run yet. (4) The donor inference rests on one MAG copy and on seven FOSC copies from one locus; the MAG’s completeness and possible chimerism have not been assessed. (5) The tblastn identities (96.7–97.7%) are lower than the DIAMOND protein identity (98.4%) for two reasons. Chained HSPs include intron-boundary edges, and the *F. tricinctum* copy carries a 1-nt deletion in a G homopolymer about 40 nt before its stop, which frameshifts the last 13 codons and truncates the protein (519 aa vs 589 aa in the four complete *F. oxysporum* gene models). DIAMOND’s 98.4% is a local identity over residues 1–506 to the best-matching strain; global identity is 95.4–96.3%. Seven of eight PacBio reads at this site agree with the assembly, but these reads are indel-prone and deeper coverage is needed to confirm the deletion; the HET domain itself (residues 23–124) is intact. (6) This candidate was the top inter-complex edge by raw similarity in the HET-domain view only (in the full-length view it ranks 494th). A graph-based scorer that we also evaluated did not rank it highly, because it cannot score recipients in clusters with fewer than three species; raw similarity should therefore be kept as a screen alongside any model-based ranking. Two other leads from the screen, older candidate transfers from Basidiomycota into the *F. solani* species complex, and a vertical control were taken through the gene-tree stage but are not reported here. (7) The stage-1 support values come from MAFFT and IQ-TREE runs with two threads, which are not deterministic (Table 2). (8) In a negative control of 56 genes not expected to have been transferred, stage 1 never produced a false pass. In simulated transfers, the chain’s sensitivity was a step at 98% protein identity, set by the stage-4 rule, so the chain detects only very recent or strongly constrained transfers. (9) Stage 5’s chained tblastn is imperfect: HSPs that run through introns lower the identities (the self-control flagged 43% of conserved genes, and 1 of 228 conserved genes was called absent from *F. avenaceum*), and a five-exon, 66-aa gene (KAH7257287.1) that is present in the panel *F. avenaceum* assembly was missed; spliced alignment (miniprot) found it.

## 5 Methods

### Genome panel and HET-domain proteins

The panel comprises 109 fungal genomes from NCBI (38 *Fusarium*, including nine *F. oxysporum* strains), with proteomes taken from the NCBI annotations and completeness assessed with BUSCO 6.0.0 (fungi_odb12, protein mode) [9, 10]. Two panel assemblies, *F. odoratissimum* NRRL 54006 (FocTR4) and *F. oxysporum* II5 (Foxii5), are the same strain; both are kept, and *F. odoratissimum* is counted as a member of the FOSC. HET-domain proteins were identified with pyhmmer [11] using the PF06985 profile and its gathering threshold (`–cut_ga`). This gave 8,927 domains in 8,904 proteins; after removing proteins shorter than 100 aa (70) and exact duplicates within a genome (19), 8,815 proteins in 89 genomes remained. Recall was checked against the 88 HET-labelled NOD-like receptors in the supplementary trees of Dyrka et al. [5]: all 88 pass the gathering threshold, and all 54 that map to panel genomes are in our set. Two sequence views were kept: the HET domain (envelope coordinates) and the full-length protein.

### Similarity screen

All-against-all DIAMOND 2.2.8 [12] searches (`blastp –ultra-sensitive`, E ≤ 10^−3^, all targets reported with `-k 0`) were run separately for both views. For each pair the best HSP was kept, similarities were normalised as 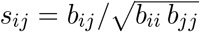 from bit scores, and the larger of the two directions was used. Edges with *s* ≥ 0.15 were analysed. Within *Fusarium*, genomes were grouped by single linkage at a species-tree patristic distance of 0.05. This gives seven clusters that largely match the species complexes; the FOSC cluster also contains the *F. fujikuroi* complex and *F. redolens*. Candidate edges join proteins from different clusters; edges between strains of one species are never candidates. The construction of the graph is described in the Supplementary Material.

### Species tree

A maximum-likelihood tree of all 109 genomes was inferred with IQ-TREE 3.1.1 [15] (LG+F+G4, unpartitioned; seed 12345) from a concatenated alignment of 318 single-copy BUSCO genes present in ≥ 85% of genomes (172,463 amino-acid columns; MAFFT L-INS-i, trimAl `-automated1`; the 318 are a subset of the 531 loci that passed alignment checks, the rest being held out). It was rooted between the two non-Dikarya genomes (Mucoromycota and Glomeromycota) and the Dikarya, and was used only after detection, to check clade membership and to compute patristic distances (Biopython).

### Gene trees

For each candidate, the closest HET-domain proteins at *s* ≥ 0.30 (up to three per species; all event proteins always kept) were collected. Domain sequences were aligned to the PF06985 HMM (match columns only); full-length sequences were aligned with MAFFT 7.526 L-INS-i [13]. Both alignments were trimmed with trimAl `-gappyout` [14]. The two views are not independent: after trimming, the full-length alignment covers mainly the N-terminal half of the protein, and 89 of its 223 columns lie in the HET domain. Trees were inferred with IQ-TREE 3.1.4 [15] with ModelFinder [16] model selection (BIC) and 1,000 ultrafast bootstrap replicates [17] (`–wbtl`, seed 12345), and midpoint-rooted for display. UFBoot support was read per bipartition of the unrooted tree. A nesting was counted as evidence only if the smallest clade containing the recipient protein had UFBoot ≥ 95 [18]. MAFFT’s iterative refinement and IQ-TREE are not deterministic when run with several threads (two here); we report the deposited run and the range over reruns. The stage-1 protocol was revised after a first run (neighbours at *s* ≥ 0.15, MAFFT alignment of domains, trimAl `-automated1`), which kept too few alignment columns. That first run gave the same placement in the domain view (*F. tricinctum* + 6 *F. oxysporum*, UFBoot 99.8); its full-length tree did not finish.

### Genome background and contamination screen

All 14,106 *F. tricinctum* proteins were searched with DIAMOND (`–sensitive`, E ≤ 10^−5^, up to 50 targets) against the combined proteomes of nine *F. oxysporum* strains, *F. avenaceum* and *F. graminearum*. For each protein and each comparator set we kept the best hit with ≥ 50% query coverage. Rules fixed beforehand: *contamination* if the HET scaffold’s median identity to *F. oxysporum* is ≥ 98% and above the genome-wide value; *block transfer* if a run of ≥ 3 consecutive genes including the HET gene is ≥ 98% to *F. oxysporum* while the rest of the scaffold is not; otherwise *single gene*. As implemented, the contamination rule compares the scaffold median with its median identity to *F. avenaceum* rather than with the genome-wide value, and the block rule does not test the rest of the scaffold; neither difference changes a verdict here. Every *F. tricinctum* scaffold was also screened for a median identity to *F. oxysporum* of ≥ 98%. A gene was called *F. oxysporum*-affine if it was ≥ 90% identical to *F. oxysporum* and ≥ 10 percentage points more identical to *F. oxysporum* than to *F. avenaceum*; a gene with no *F. avenaceum* hit counted as *F. oxysporum*-affine if it was ≥ 90% identical to *F. oxysporum*. Lineage-specific chromosomes of *F. oxysporum* f. sp. *lycopersici* 4287 were taken as the four entirely lineage-specific chromosomes 3, 6, 14 and 15 [6]; the lineage-specific parts of chromosomes 1 and 2 and the unanchored scaffolds were not counted.

### Sister-species control

*F. tricinctum* proteins whose best *F. oxysporum* hit is ≥ 98% identical were split into those also ≥ 95% identical to *F. avenaceum* (conserved) and those absent from, or < 70% identical to, *F. avenaceum* (donor-like, sister-absent). The 11 sister-absent genes were then searched in 15 further *F. avenaceum* genomes (tblastn with the stage-5 rules) and, with spliced alignment (miniprot [43]), in all 16 *F. avenaceum* assemblies. A gene was reclassified as not donor-like if any *F. avenaceum* genome carried a close copy (≥ 90%); the result under the stage-4 rule itself (any copy ≥ 70%) is also reported.

### Assembly absence check

Genome assemblies of the 38 *Fusarium* genomes were downloaded from NCBI. Each was searched with tblastn (BLAST+ 2.16 [19]; E ≤ 10^−5^, up to 50 targets) using the HET protein and the four other sister-absent scaffold-9 proteins as queries. HSPs on the same contig and strand with gaps ≤ 20 kb were chained into loci; per locus we computed the summed bit score, query coverage (union of HSP query ranges) and length-weighted identity, and kept the locus with the highest summed bit score. Calls were fixed beforehand: close copy, ≥ 90% identity over ≥ 50% of the query; absent, best locus < 70%; related copy, in between. A query whose self-search against the *F. tricinctum* assembly did not return a close copy was marked unreliable. The same procedure was applied to 24 further public assemblies of the *F. tricinctum* complex and to the MAG GCA_027247665.1 (Supplementary Table S1).

### Synonymous divergence

Coding sequences were taken from the NCBI annotations and assemblies (translations checked against the proteomes) for every *F. tricinctum* protein and its best *F. oxysporum* hit (stage 3), for every *F. tricinctum* protein and its best *F. avenaceum* hit, for the best Fo47 hits of a random sample of 1,500 Fol 4287 proteins (1,490 pairs), and for the HET gene and each *F. oxysporum* copy. Pairs were aligned as proteins and back-translated to codons, and *d*_*S*_ was computed with the Nei–Gojobori method and a Jukes–Cantor correction [36]. Pairs with fewer than 51 aligned codons were dropped, and pairs with *p*_*S*_ ≥ 0.75 were treated as saturated. With other estimators (LWL85 [37], YN00 [38], maximum likelihood [39] and fourfold-degenerate sites), with reciprocal-best-hit backgrounds and on the in-frame part of the HET gene (codons 1–506), the HET copies have *d*_*S*_ 0.05–0.12; this is below 99.5–100% of *F. tricinctum*–*F. oxysporum* backgrounds and 86–97% of *F. tricinctum*–*F. avenaceum* backgrounds.

### Nucleotide gene tree and topology tests

For the HET gene and its close and related copies, the region homologous to the *F. tricinctum* gene (from the start codon to the position of the *F. tricinctum* stop codon, intron included) was taken from each assembly at the locus found by spliced alignment of the *F. tricinctum* protein (miniprot), aligned with MAFFT L-INS-i (one thread) and analysed with IQ-TREE 3.1.4 (ModelFinder, BIC; 1,000 UFBoot with `-bnni`; 1,000 SH-aLRT replicates [41]; seed 12345). Two sets of sequences were used. The first holds the *F. tricinctum* copy, the seven FOSC copies and the MAG copy, with the *F. proliferatum* and *F. flagelliforme* copies as outgroups (11 sequences, 1,614 columns, HKY+G4). The second adds five further related copies (16 sequences, 1,629 columns, K2P+I; Fig. 6). Support values were read per bipartition. Trees constrained to each alternative placement were compared with approximately unbiased (AU) tests [40] (10,000 RELL replicates). Protein-level AU tests used a 15-sequence alignment of the close and related copies under three models.

### Mobile-element context and controls

The regions around the HET gene in *F. tricinctum* and around the FOSC copies were searched for transposon domains and the *Starship* captain domain (DUF3435) in annotated proteins and six-frame open reading frames (pyhmmer, Pfam), and repeats were found by aligning the regions to their own genomes (dc-megablast); the strain-specific island was defined by synteny with the MsR-QD66 and “*F. acuminatum*” 1A genomes. As controls, 56 genes not expected to have been transferred (conserved genes and typical single-copy genes) were run through a fast version of stage 1 and through stage 5, and 20 Fo47 genes without *F. tricinctum* or *F. avenaceum* homologs were inserted into a copy of the *F. tricinctum* genome at 0–20% nucleotide divergence and run through stages 3–5.

### Read support and genome alignments

The first 300,000 PacBio reads of the sequenced strain (SRA run SRR7777439) were mapped to its assembly with minimap2 [42] (`map-pb`; MAPQ ≥ 20) to compute depth and reads spanning the HET gene. Scaffold 9 and core scaffold 1 were aligned to other assemblies with minimap2 (`asm20`) to measure shared sequence and genome-wide identity, and the FOSC loci were placed on Fol 4287 chromosomes with minimap2 (`asm10`).

## Data availability

All genomes, proteomes and annotations are public in NCBI GenBank/RefSeq under the accessions listed in Supplementary Table S1 (e.g. *F. tricinctum* GCA_020744515.1, *F. avenaceum* GCA_020744115.1, *F. oxysporum* f. sp. *lycopersici* 4287 GCF_000149955.1, MAG GCA_027247665.1). PacBio reads of *F. tricinctum* MPI-SDFR-AT-0068 are in the SRA (SRR7777438, SRR7777439).

