## Supplemental Material for "A staged evidence chain identifies a recent horizontal transfer of a HET-domain gene into a *Fusarium tricinctum* strain from a lineage related to the *Fusarium oxysporum* species complex"

### Supplementary Material:
### Building the HET-domain similarity graph

#### 1 Introduction

The HET-domain similarity graph has **8,815 nodes** and **18,552,568 undirected weighted edges**, or **12,397,947 edges at the analysis cut** $s \geq 0.15$ (density 0.319). Each node is one HET-domain protein (Pfam PF06985, gathering threshold) from 89 of 109 fungal genomes. Each edge joins two proteins whose excised HET domains align in an all-vs-all DIAMOND blastp search. The weight is the self-score-normalised bit score $s = b_{ij}/\sqrt{b_{ii}\, b_{jj}}$. The genome panel, proteomes, BUSCO scores, assembly metadata, HMM and species tree were generated. The HMM search, the QC, the DIAMOND run, the normalisation and all node metadata were redone in this project. Node metadata (NCBI taxonomy, gene, domain coordinates, copy number, paralog family, track groups and genomic context) sits in tables keyed by `node_index`, not inside the matrix file.

#### 2 Data sources

Every sequence in the graph traces back to a public NCBI genome annotation. 79 of the 109 genomes are RefSeq (GCF) and 30 are GenBank (GCA). Table 1 lists the inputs; the genomes and their accessions are listed in Supplementary Table S1 (file `TableS1_genomes.tsv`).

#### 3 Nodes

A node is one HET-domain protein that passes QC: 8,815 nodes in 89 genomes (`build_graph.pyhet-extract`).

1. **Search.** pyhmmer 0.12.3 `hmmsearch` with PF06985 and `bit_cutoffs="gathering"`, i.e. HMMER's `--cut_ga`, over all 109 proteomes. A protein is kept if its sequence score is $\geq 22.5$ and at least one domain scores $\geq 22.5$. Non-letter characters such as `*` were masked to X for the search only. Result: 8,927 GA-passing domains in 8,904 proteins from 89 genomes. 20 genomes have no HET protein at GA.

2. **QC** (`configs/pilot.yaml`, `data.het_extraction`), applied in this order:

   - length $\geq 100$ aa: 70 proteins failed;
   - ambiguous residues (X, B, Z, J, U, O, *) $\leq 10\%$: none failed;
   - exact duplicate sequence within the same genome: 19 removed, keeping the first protein ID in sorted order. Alleles and paralogs that differ by one residue stay.

3. **Two sequence views of each node.** Both FASTA files hold the same IDs in the same order.

**Table 1: Inputs to the graph**

| Input | Content | Original source |
|---|---|---|
| Genome panel | 109 genomes: 91 Ascomycota, 16 Basidiomycota, 1 Glomeromycota, 1 Mucoromycota; 38 *Fusarium*, incl. 9 *F. oxysporum* strains (*F. odoratissimum* NRRL 54006, also in the *F. oxysporum* complex, is the same strain as *F. oxysporum* II5); 100 NCBI species, 60 genera, 49 families | NCBI Assembly, previously chosen |
| Proteomes | One protein per gene: CDS grouped by gene in the GFF3, longest product kept | NCBI protein.faa + GFF3 by accession |
| Assembly metadata | Organism name, NCBI tax_id per accession | NCBI Datasets assembly reports |
| Taxonomy | Phylum, class, order, family, genus, species per tax_id | NCBI Datasets taxonomy API, fetched once by this project and cached |
| BUSCO | Complete and duplicated % per genome | BUSCO 6.0.0, fungi_odb12, proteins mode |
| HET HMM | Pfam PF06985.18 (HET), 148 match states, GA 22.5 / 22.5 bits | Pfam |
| Species tree | IQ-TREE 3.1.1 ML tree (LG+F+G4) on 318 concatenated single-copy BUSCO genes (172,463 amino-acid columns), 109 tips, rooted between the two non-Dikarya genomes (Mucoromycota, Glomeromycota) and the Dikarya | |
| GFF3 + sequence reports | Genomic context of each node | NCBI, fetched for this project for all 109 genomes |

- het_domain.faa (primary, used for this graph): the envelope (env_from–env_to) of the protein's best-scoring GA domain. The 23 proteins with more than one HET domain contribute only that best domain. The median length is 148 aa (range 45–247).
- het_full.faa: the whole protein. It is used for the companion full-length graph.

4. **Node ID and order.** IDs are <shortname>__<proteinaccession>, e.g. Fol_4287__XP_018258429.1. s1_diamond/nodes.tsv fixes the integer order 0–8,814. Row and column $k$ of the matrix, and node_index $= k$ in the node metadata tables (column index in nodes.tsv), all refer to the same protein.

By phylum, the nodes are 8,566 Ascomycota and 249 Basidiomycota. 5,473 (62%) are *Fusarium*. Copy number ranges from 1 to 499 per genome (median 93 over the 89 genomes with HET proteins). *Hyaloscypha* has 499 nodes, and *Trametes versicolor*, with 172, has the most among the Basidiomycota.

### 4 Edges

An edge joins two nodes if DIAMOND reports any alignment between their HET domains at $E \leq 10^{-3}$. Its weight $s$ is the normalised bit score below (build_graph.pydiamond).

1. **All-vs-all search.** DIAMOND 2.2.8 (bioconda, aarch64): makedb on het_domain.faa, then blastp of the same file against itself with --ultra-sensitive-e1e-3-k0. -k0 reports every

**Table 2: Items from Previous Project**

| Items | Used for |
|---|---|
| Panel definition | List of 109 genomes (shortname, accession, organism) |
| Proteomes | Sequences searched for HET; `gene=` header tag gives the gene name |
| Assembly metadata | tax_id per accession, used to fetch taxonomy |
| BUSCO summary | `busco_complete`, `busco_duplicated` in `species.tsv` |
| PF06985 HMM | HET search profile and its GA cut-offs |
| Species tree | Track B groups only, after detection; never used to build edges |

target, so the all-vs-all is complete rather than truncated at the top hits. This gave 35,735,190 HSPs.

2. **Self-scores.** $b_{ii}$ is the bit score of node $i$'s hit to itself. All 8,815 nodes had one, so none needed the fallback rerun with `--masking0`.

3. **Best HSP per ordered pair.** For each (query $i$, subject $j$), $i \neq j$, the HSP with the highest bit score is kept: 35,726,375 directed pairs, saved to `pairs_best_hsp.tsv.gz`.

4. **Normalise.** Each directed pair gets

$$s_{ij} = \frac{b_{ij}}{\sqrt{b_{ii}\, b_{jj}}}, \tag{1}$$

where $b_{ij}$ is the pair's best-HSP bit score. $s$ runs from 0 to 1, where 1 means as similar as each domain is to itself. Five values above 1 were clipped to 1.

5. **Symmetrise.** The undirected weight is $s = \max(s_{ij}, s_{ji})$, and the diagonal is set to 0. If only one direction was reported, that value is used.

6. **Store.** A symmetric SciPy CSR matrix with 18,552,568 undirected edges, saved as `S.npz`. No $s$ threshold is applied at this stage.

7. **Threshold downstream.** All analyses use $s_{\min} = 0.15$, with 0.20 and 0.25 as consistency checks. Cuts below about 0.13 increasingly coincide with the E-value filter, which already sets a per-pair floor of $s \approx 0.06$–$0.13$ (5th–95th percentile; median 0.094; `pipeline/checks/floor_check.py`; Fig. 1); a cut at 0.10 would remove only 0.5% of edges.

No taxonomy, tree or metadata is used to make or weight an edge. The graph is built from sequence alone. Groups and the species tree are applied only after detection, to choose which edges count as candidates.

### 5 Metadata

The matrix `S.npz` holds only the edge weights. Every attribute lives in a TSV row keyed by `node_index` (nodes) or `shortname` (genomes), so the graph and its labels join on one integer index. Edges carry one stored attribute, $s$. Edge-level labels such as same-species, between-group or candidate are derived from the two endpoint nodes' metadata when needed (Table 4). Every node carries the same 23 attributes, in the column order of Table 3. Table 4 gives where each attribute comes from, plus the genome-, pair- and context-level attributes stored in other files.

**Table 3: The 23 node attributes** (columns of `het_proteins.tsv`, in file order).

| Group | Columns |
|---|---|
| Identity | `node_index`, `protein_id`, `shortname` |
| Genome | `accession`, `organism`, `tax_id` |
| Taxonomy | `phylum`, `class`, `order`, `family`, `genus`, `species` |
| Gene | `gene` |
| HET domain | `protein_len`, `n_domains`, `best_domain_score`, `env_from`, `env_to`, `domain_len`, `hmm_coverage` |
| Copy number | `copy_number` |
| Paralog family | `within_taxon_family_size`, `within_taxon_family` |

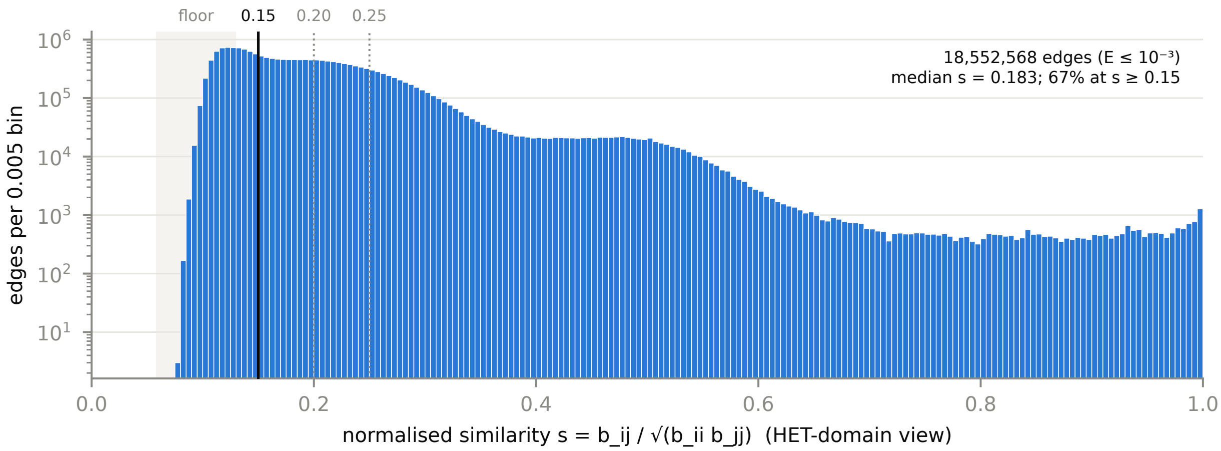


**Figure 1: Distribution of edge weights in the HET-domain graph.** All 18,552,568 edges ($E \leq 10^{-3}$) in bins of 0.005; the $y$ axis is logarithmic. The shaded band is the 5th–95th percentile of the per-pair floor that the E-value filter alone imposes on $s$; the solid line is the analysis cut ($s \geq 0.15$) and the dotted lines the consistency checks (0.20, 0.25).

#### 6 Size and density

The graph is very dense: at the analysis cut $s \geq 0.15$ it keeps **12,397,947 of 38,847,705 possible edges (31.9%)**, and the median node has 3,228 neighbours. Density $= 2m/(n(n-1))$ with $n = 8{,}815$ (Table 5).

Edge weights are low in bulk: over all 18.55M edges the median $s$ is 0.183 (5th–95th percentile 0.112–0.350; Fig. 1). Among edges between different species at $s \geq 0.15$, the median $s$ is about 0.22 at every taxonomic distance, from same genus to different phyla (0.21–0.24). Only the upper tail falls, overall, with distance: the 99.9th percentile drops from 0.94 within a genus to 0.56 between phyla. 98% of edges at $s \geq 0.15$ join different genomes.

Within the two analysis tracks at $s \geq 0.15$, track A (35 NCBI-family groups among the genomes with HET proteins) has 5,754,294 candidate edges between families with at least one *Fusarium* end. Track B (7 *Fusarium* clades) has 3,526,034 candidate edges between clades. The companion full-length graph has 16.25M edges in total and 1.0M at $s \geq 0.15$.

### 7 Diagram: one corner of the graph

Figure 2 shows the neighbourhood of the *F. oxysporum* → *F. tricinctum* lead: the *F. tricinctum* node, its three closest *F. oxysporum* matches (three further *F. oxysporum* copies, $s = 0.88$–$0.97$, are not drawn), the closest nodes of two other *Fusarium* species (*F. flagelliforme* and *F. proliferatum*), the closest node from its sister species *F. avenaceum*, the closest *F. graminearum* node and the closest non-*Fusarium* node. In the real graph every one of these 9 nodes has more than 1,300 neighbours at $s \geq 0.15$, so the drawing shows 36 of their about 12,800 edges. The two boxes show how the data attaches: node attributes live in `het_proteins.tsv` (and `s1_gff`), the edge stores only $s$, and candidate labels are computed from the endpoints' groups when a track uses them.

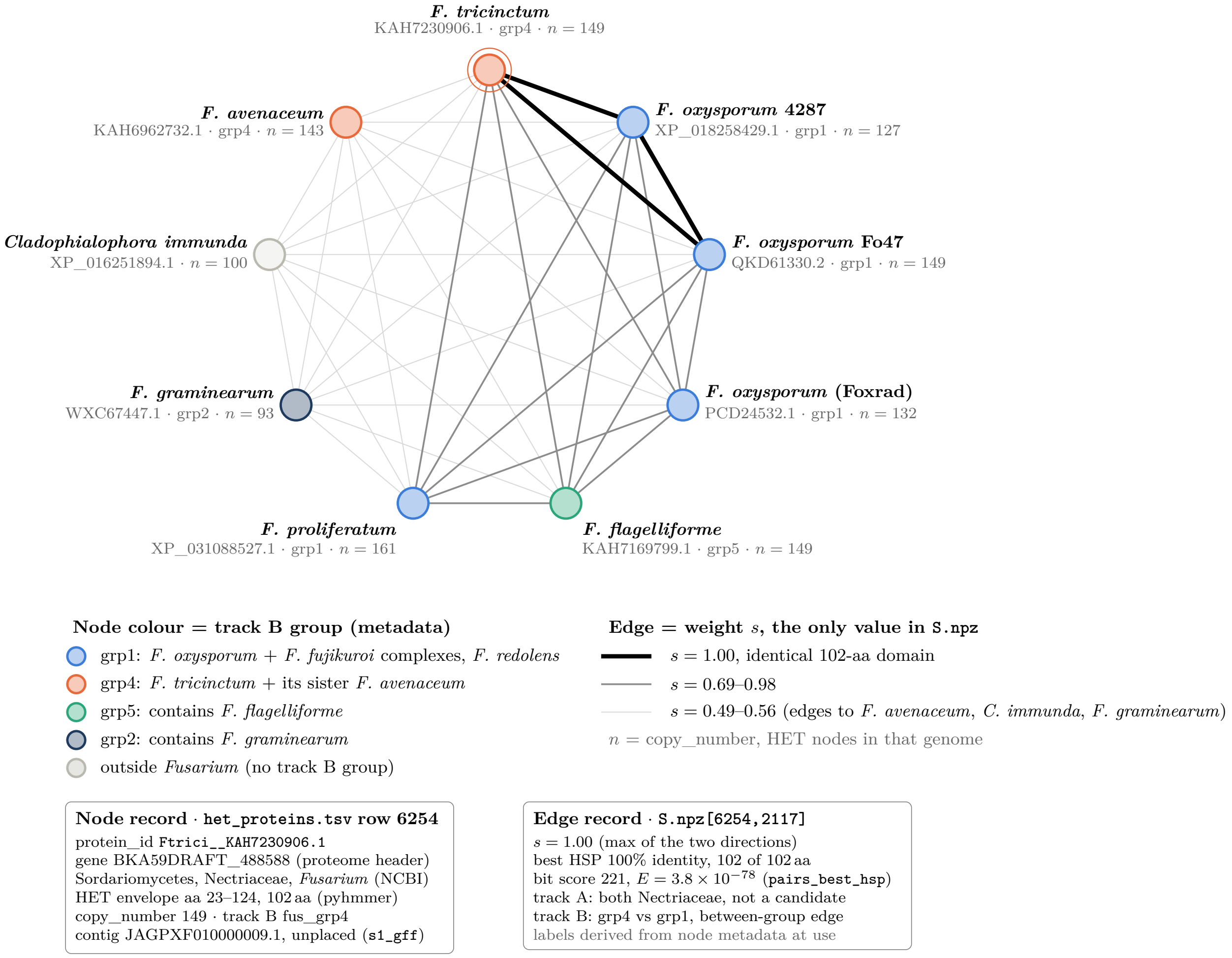


**Figure 2: Identical HET domains join *F. tricinctum* to *F. oxysporum*.** Nine of the 8,815 nodes and all 36 edges among them (het_domain view, real $s$ values). Each of these nodes has 1,349–1,707 neighbours at $s \geq 0.15$ in the full graph.

Software: pyhmmer 0.12.3, DIAMOND 2.2.8, NumPy/SciPy sparse, pandas, Biopython (track B tree distances), NCBI Datasets API and CLI 18.38.0. Load the graph with `scipy.sparse.load_npz("S.npz")` and join rows to `het_proteins.tsv` on `node_index`.

**Table 4: Node, genome and edge attributes**

| Attribute | Level | Files stored in | Source |
|---|---|---|---|
| `node_index`, `protein_id`, `shortname` | node | `nodes.tsv`, `het_proteins.tsv` | Node order from `het_full.faa`; selected IDs |
| `accession`, `organism`, `tax_id` | node, genome | `het_proteins.tsv`, `species.tsv` | YAML + assembly JSONL (NCBI) |
| `phylum` ... `species` (6 ranks) | node, genome | `het_proteins.tsv`, `species.tsv` | NCBI Datasets taxonomy API by tax_id; raw replies cached in `ncbi_taxonomy_reports.json`. Two genomes (*Yarrowia lipolytica*, *Mycosarcoma maydis*) have no NCBI family |
| `gene` | node | `het_proteins.tsv` | `gene=` from the NCBI GFF3 |
| `protein_len`, `n_domains`, `best_domain_score`, `env_from`, `env_to`, `domain_len`, `hmm_coverage` | node | `het_proteins.tsv` (also `het_proteins_s1.tsv`) | This project's pyhmmer PF06985 search |
| `copy_number` ($n_i$) | node | `het_proteins.tsv` | Count of HET nodes in the node's genome |
| `within_taxon_family`, `within_taxon_family_size` | node | `het_proteins.tsv` | Single linkage within each genome over full-length DIAMOND best-HSP pairs with identity ≥ 80% covering ≥ 80% of both proteins: 8,786 families, 8,763 singletons, largest 5 |
| `busco_complete`, `busco_duplicated` | genome | `species.tsv` | BUSCO summary |
| `n_het_nodes`, `n_panel_same_species`, `n_within_taxon_families` | genome | `species.tsv` | Computed here |
| `self_bitscore` ($b_{ii}$) | node | `self_scores.tsv` | DIAMOND self-hit |
| `pident`, coordinates, `evalue`, `bitscore`, lengths | directed pair | `pairs_best_hsp.tsv.gz` | DIAMOND best HSP |
| Track A group (NCBI family) | genome | `genome_groups.tsv` | NCBI family; a genome with no family is its own group (35 groups among the 89 genomes with HET proteins; the file lists all 109 genomes, in 51 groups) |
| Track B group (*Fusarium* clade) | genome | `genome_groups.tsv` | Single linkage at patristic distance ≤ 0.05 on the species tree, giving 7 groups; group 1 holds the *F. oxysporum* and *F. fujikuroi* complexes and *F. redolens* |
| Contig, role, chromosome, coordinates, strand, locus tag, gene rank, distance to contig end, product | node | `het_gene_context.tsv`; ±10 flanking genes in `het_flanks.tsv` | NCBI GFF3 + assembly sequence report, re-fetched here. All 8,815 nodes placed. Used for evaluation only |

**Table 5: Graph size and density at each cut** (het_domain view, $n$ = 8,815 nodes). Inter-genome edges join nodes from different genomes.

| Cut | Edges | Density | Mean degree | Median degree | Max degree | Isolated nodes | Inter-genome edges | Components |
|---|---|---|---|---|---|---|---|---|
| none ($E \leq 10^{-3}$) | 18,552,568 | 0.478 | 4,209 | 4,507 | 7,956 | 0 (min deg. 24) | 18,181,477 | 1 |
| $s \geq 0.15$ (primary) | 12,397,947 | 0.319 | 2,813 | 3,228 | 6,480 | 1 | 12,142,559 | 2 (8,814 + 1) |
| $s \geq 0.20$ | 7,737,711 | 0.199 | 1,756 | 1,845 | 3,715 | 8 | 7,571,089 | 11 |
| $s \geq 0.25$ | 3,833,424 | 0.099 | 870 | 848 | 2,939 | 35 | 3,745,508 | 48 |
